# Effect of eribulin on angiogenesis and endothelial adhesion molecules

**DOI:** 10.1101/2021.10.28.466220

**Authors:** Andrea Abbona, Antonella Falletta, Matteo Paccagnella, Simonetta Astigiano, Stefania Martini, Nerina Denaro, Fiorella Ruatta, Ottavia Barbieri, Marco Merlano, Ornella Garrone

**Affiliations:** Lab of translational oncology, ARCO foundation, S. Croce & Carle Teaching Hospital, Cuneo, Italy; IRCCS Ospedale Policlinico San Martino, Genova, Italy; Oncology Dept, S. Croce & Carle Teaching Hospital, Cuneo, Italy; Dept of Experimental Medicine, University of Genova, Genova, Italy; Oncology Dept, Candiolo Cancer Institute, FPO-IRCCS, Candiolo (Torino), Italy; Breast Unit, Oncology Dept, S. Croce & Carle Teaching Hospital, Cuneo, Italy

**Keywords:** eribulin, TGF-β, EndMT, neo-angiogenesis, adhesion molecules

## Abstract

Tumor vasculature is an important component of the tumor microenvironment and deeply affect anticancer immune response. Eribulin is a non taxane inhibitor of the mitotic spindle. However, off-target effect interfering with the tumor vasculature have been reported. The mechanisms responsible of this effect is not clear.

We designed an *in vitro* study to investigate the effect of eribulin on neo-angiogenesis and on the adhesion molecules ICAM-1 and VCAM-1, with or without TGF-β. We also investigated the effects of paclitaxel and vinorelbine in the same experimental conditions.

Eribulin was able to up-regulate the epithelial markers VE-cadherin and CD-31 in the HUVEC and tube formation in HUVEC cultured in Matrigel. The drug effectively arrested tube formation even in presence of TGF-β. Eribulin counteracted the TGF-β induced change in cell shape from the endothelial cobblestone-like morphology to an elongated spindle-shaped morphology that is characteristic of EndMT.

We also observed that eribulin is able to upregulate ICAM-1 and to counteract its downregulation induced by TGF-β.

In this study, eribulin was able to inhibit the vasculature remodeling and the downregulation of ICAM-1 induced by TGF-β. These effects might have important therapeutic consequence if the drug will be administered with immunotherapy.

## 1 Introduction

Microtubules are important target of cancer chemotherapy. Eribulin, as well as vinorelbine, paclitaxel and other agents, directly targets microtubules. Unlike these drugs, eribulin inhibits microtubules growth causing nonproductive tubulin aggregates without any effect on microtubules shortening. This leads to arrest the formation of the mitotic spindle, irreversible mitotic block at G2-M phase and apoptosis (reviewed in 1).

Eribulin is approved for the treatment of metastatic breast cancer by EMA after at least one previous line of therapy, and by FDA for the treatment of metastatic breast cancer after at least two previous lines of therapy.

In addition to the main mechanism of action on microtubules, eribulin shows off-target effects able to interact with the tumor microenvironment (TME).

In particular, eribulin is able to interfere with neo-angiogenesis (2), the expression of TGF-β and of other cytokines both in experimental models and in humans (3, 4).

TGF-β interferes with the tumor microenvironment leading to an immunosuppressive environment and strongly contributes to T-cells exclusion through many mechanisms such as the induction of reactive stroma and neo-angiogenesis (5). Therefore, the modulation of TGF-β by eribulin might favor the homing of effector T cells in excluded tumors.

Many human breast cancers are immune excluded tumor, in particular triple negative breast cancer (6) and, as such, are characterized by strong TGF-β signal, neo-angiogenesis and reactive stroma (5).

These cancer characteristics may have contributed to the success of eribulin in this disease.

The deep knowledge of the off-target effects of eribulin may support its use in combination with immune therapy.

We designed the following *in vitro* study to investigate the effect of eribulin on the human vascular endothelial cell (HUVEC) in terms of endothelial-mesenchymal transition (EndMT) and expression of endothelial adhesion molecules (ICAM-1 and VCAM-1).

## 2. Materials & Methods

### 2.1 Study design

Eribulin effect was evaluated on HUVEC with or without previous exposure to TGF-β.

We also tested the effects of navelbine and paclitaxel in the same experimental conditions and and compared the results with those achieved with eribulin and TGF-β alone.

### 2.2 Cell culture

The Human Umbilical Vein Endothelial Cells (HUVEC) were purchased from Lonza (Walkersville, MD, USA; batch 0000440546 and 0000442486) and cultured in EBMTM-2 Basal Medium (Lonza, Walkersville, MD, USA) with EGM-2MV Single Quots (Lonza, Walkersville, MD, USA) in 25 cm^2^ gelatinized cell culture flasks at 37 °C, 5% CO_2_. All the experiments were performed using cells at passage number between 5 and 12.

### 2.3 Drugs

TGF-β, TNF-α and IL-1β were purchased from Life Technologies Europe (Bleiswijk, The Netherlands), reconstituted and stored according to manufacturer’s recommendations. Eribulin (Halaven®, EISAI, Tokyo, Japan), paclitaxel (Taxol® Bristol-Meyers Squibb, NY, USA) and vinorelbine (Navelbine 50® Pierre Fabre, Paris, France) were provided by the Santa Croce and Carle Hospital pharmacy and Ospedale Policlinico IRCCS S. Martino, Genova, and kept at 4 °C for no longer than 7 days. For treatments the drugs were freshly diluted in cell medium at the chosen concentrations.

### 2.4 HUVEC morphology and tube formation assay on matrigel

HUVEC were seeded in 25 cm^2^ gelatinized cell culture flasks and treated for 24 hours with 1 nM (2xIC_50_) (7) eribulin, or 10 ng/ml TGF-β (8), or the combination of the two. Images were acquired using inverted Leika DR IRB microscope associated to a Nikon camera.

For tube formation on matrigel, 24 well plates were coated with 200 μl of growth factors depleted matrigel (BD, Franklin Lakes, NJ, USA) and incubated 30 min at 37 °C. HUVEC were then seeded at 7×10^4^ cells/well density in 500 μl of growth medium and monitored for network formation up to 24 hours. Drugs were added during seeding, either alone or in combination, at the final concentration of 10 ng/ml TGF-β and 1 nM eribulin. Experiments were repeated 4 times. Images were acquired after 4 hours using an Olympus stereo-microscope (Olympus Corp. Tokyo, Japan).

### 2.5 RNA isolation and quantitative real-time (qRT-PCR)

To measure the 2 epithelial markers (VE-cadherin and CD31) and the 3 mesenchymal markers (Vimentin, Snail and αSMA) in the first set of experiments HUVEC were seeded in 25 cm^2^ flasks and, at confluence, treated with either 1.0 nM eribulin, 0.8 nM paclitaxel, 10 nM vinorelbine or medium for 4 hours. In the second set of experiments cells were pre-treated with 10 ng/mL TGF-β for 4 hours, subsequently cells were washed and treated with medium, eribulin, paclitaxel, or vinorelbine at the above-mentioned concentrations for 4 more hours. Each experiment was performed in triplicate.

Total RNA was extracted from each cell sample using RecoverAll ™ Total Nucleic Acid Isolation Kit (Ambion Inc, Austin, TX, USA) according to the manufacturer’s instructions. Concentration and purity of the RNA samples were tested using NanoDrop ND 1000 (Thermo Fisher Scientific, Wilmington, Delaware, USA) at 260/280 nm. Total RNA sample was reverse transcribed to cDNA using High Capacity cDNA Reverse Transcription Kit (Applied Biosystems, Foster City, CA, USA) according to manufacturer’s instructions, using Applied Biosistems Veriti 96 well Thermal Cycler 1000 (Thermo Fisher Scientific, Wilmington, Delaware, USA). The reaction conditions were: 10 minutes at 25 °C, followed by 2 hours at 37 °C and 5 minutes at 85 °C. The PCR assay was performed in duplicate in a total volume of 25 μl, using 12.5 μl SYBR™ green chemistry (SYBR™green master mix, Applied Biosystems, Foster City, CA, USA) 4 μl of c-DNA and 1.25 μl of 50 uM each primer for 40 cycles. Supplementary Table 1 shows the genes analyzed and the primers used. mRNA fold change was calculated using the 2^-ΔΔCt^ method (9) where GAPDH was used as control gene.

### 2.6 Flow cytometry analysis

The effect of eribulin on the expression of the membrane proteins ICAM-1, and VCAM-1 was analyzed by flow cytometry. Endothelial cells were identified staining with CD31 antibody. HUVEC were seeded overnight onto 6 well plates coated with 0,1% gelatin, then treated for 24 hours with 1 or 10 ng/ml TGF-β, for VCAM-1 or ICAM-1 respectively, 10 ng/ml TNF-α (7); 5 nM eribulin (10xIC_50_), and combinations thereof. Cells were then detached by a brief incubation in 0,012% trypsin/ 0,1 mM EDTA and stained with the following monoclonal antibodies: PE mouse anti-human ICAM-1, PE mouse anti-human VCAM-1 or Alexa Fluor 647 mouse anti-human CD31 (all antibodies were from BD, Franklin Lakes, NJ, USA). Samples were analyzed on Gallios Flow Cytometer (Beckman Coulter, Brea, CA USA), data were processed with FlowJo software (TreeStar, Ashland, OR, USA).

### 2.7 Statistical analysis

The statistical analyses were conducted using GraphPad Prism 5. Statistical significance for transcript analyses was determined by non-parametric Mann-Whitney *U* test. Considering the number of comparisons planned, we adjusted for multiple testing using the Bonferroni correction. For flow cytometry analysis, statistical significance was determined by the paired Student t test. Data are expressed as mean ± SD.

To analyze the significance in molecular markers expression changes induced by the treatments, the data of the individual expressions were collected in a *ratio* (E/M) calculated between the sum of the *ratio* of the 2 epithelial markers (VE-cadherin and CD31 in the numerator) and the sum of the *ratio* of the 3 mesenchymal markers (Vimentin, Snail and αSMA in the denominator). The calibrant used were the untreated HUVEC (CTR). If a value of ΔΔCT is > 0, corresponding to the condition ΔCT (CTR)>ΔCT (test) and replacing its value in the expression of the *ratio* (2^-ΔΔCT^) a *ratio* <1 was obtained, the under-expression index of the mRNA of interest in the treated sample, compared to CTR. On the contrary, a value of ΔΔCT <0, that is ΔCT (CTR) <ΔCT (test), a *ratio* >1 was obtained, which corresponds to an increase in expression of the mRNA in the treated sample compared to CTR.

## 3 Results

### 3.1 Eribulin affects morphology and tube formation of HUVEC

To test whether eribulin influences EndMT and vascular remodeling, we first checked the morphological effect of the drug on HUVEC monoleyers, either alone or in the presence of TGF-β which is known to induce Epithelial Mesenchymal Transition (EMT) and EndMT (10).

Eribulin was administered for 24 hours at the concentration of 1 nM.

TGF-β was given at 10 ng/ml, accordingly to literature. TGF-β induced a change in cell shape from the endothelial cobblestone-like morphology to an elongated spindle-shaped morphology that is characteristic of EndMT. Eribulin did not induce HUVEC EndMT but effectively counteracted the effect of TGF-β (Fig. 1A).

**Figure 1.**
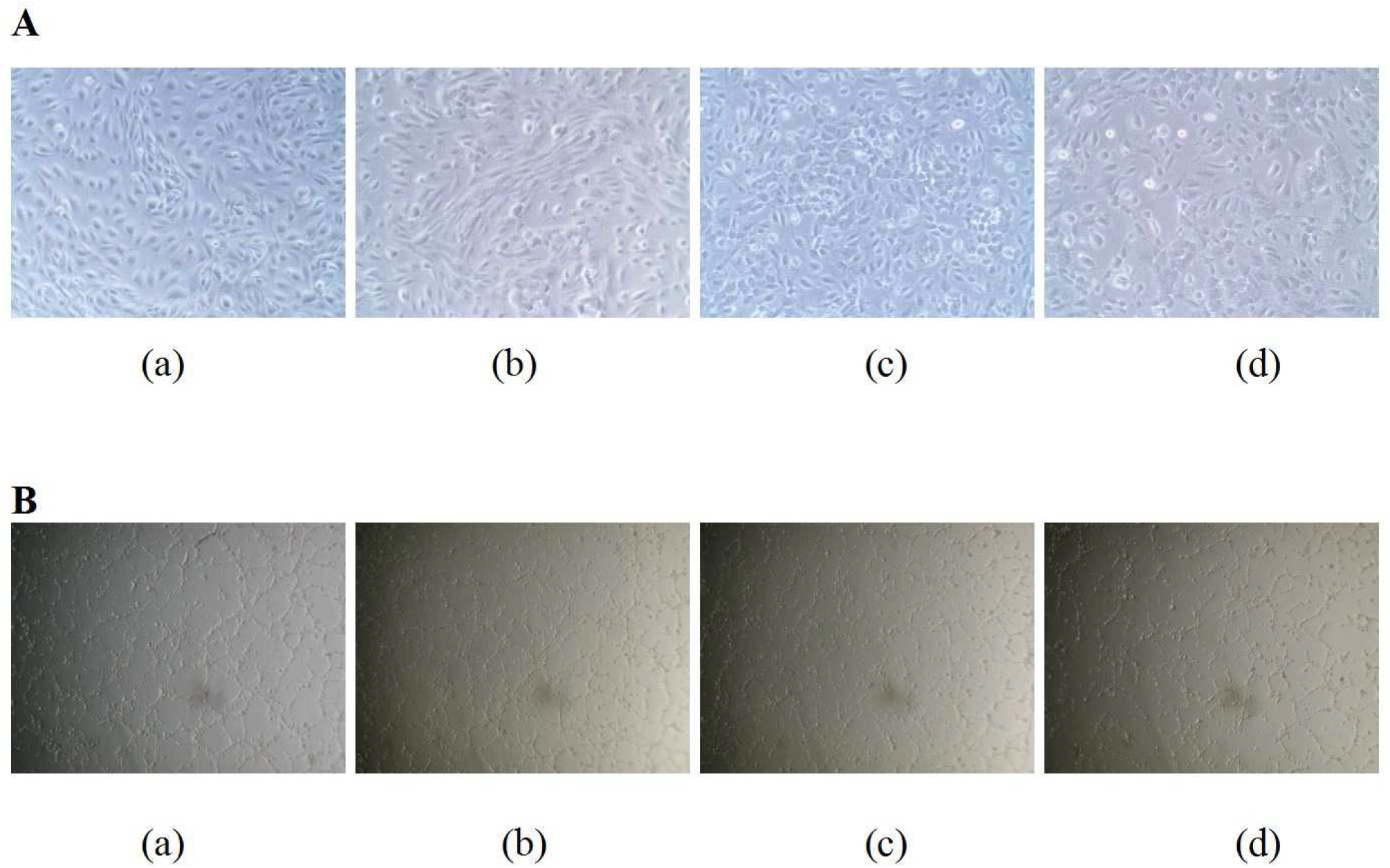
Morphogenesis of HUVEC incubated with eribulin and/or TGFβ. (A) Morphology was studied on HUVEC incubated in gelatinized flasks for 24 hrs with medium (a), 10 ng/ml TGFβ (b), 1 nM eribulin (c), or TGF-β + eribulin (d). Images were taken at x500 magnification. (B) Tube formation was studied on HUVEC incubated on matrigel-coated plates with medium (a), 10 ng/ml TGF-β (b), 5 nM eribulin (c), TGF-β + eribulin (d). Images were taken at ×750 magnification 4 hours after seeding.

Eribulin has been reported to alter tumor microenvironment also through vascular remodeling (2). Thus, we tested its influence on the ability of HUVEC to organize a reticulum after seeding on matrigel (11). In cells incubated with medium or TGF-β, a well-organized tubule network was detected 4 hours after seeding (Fig. 1B, a and b). On the contrary, in cells treated with eribulin the reticulum was not completely organized, displaying shorter and disjoined ramifications (Fig. 1B, a and c). The same phenotype was detected when eribulin was added together with TGF-β (Fig. 1B, c and d), indicating that also under these conditions, eribulin was able to overcome the TGF-β effect.

### 3.2 Eribulin modulates the expression of EndMT markers in HUVEC

To better elucidate the effect of eribulin on EndMT and vascular remodeling, we evaluated the variation of molecular markers involved in the EndMT, namely Vimentin, Snail, αSMA, CD31 and VE-cadherin (12) by qPCR.

Moreover, to check whether the effect of eribulin on EndMT was similar to other inhibitors of the mitotic spindle, we tested the effect of vinorelbine (10 nM) and paclitaxel (0.8 nM) on the same molecular markers (13, 14). Finally, TGF-β was also tested.

Eribulin did not affect the transcription of the mesenchymal markers compared to CTR, while vinorelbine upregulated Snail and paclitaxel downregulated transcription of both Vimentine and Snail.

The transcription of CD31 and VE-cadherin was significantly enhanced by all the drugs. However, eribulin had the strongest effect on the increase of VE-cadherin mRNA, while vinorelbine effect was highest on the increase of CD31 (Fig. 2A).

**Figure 2.**
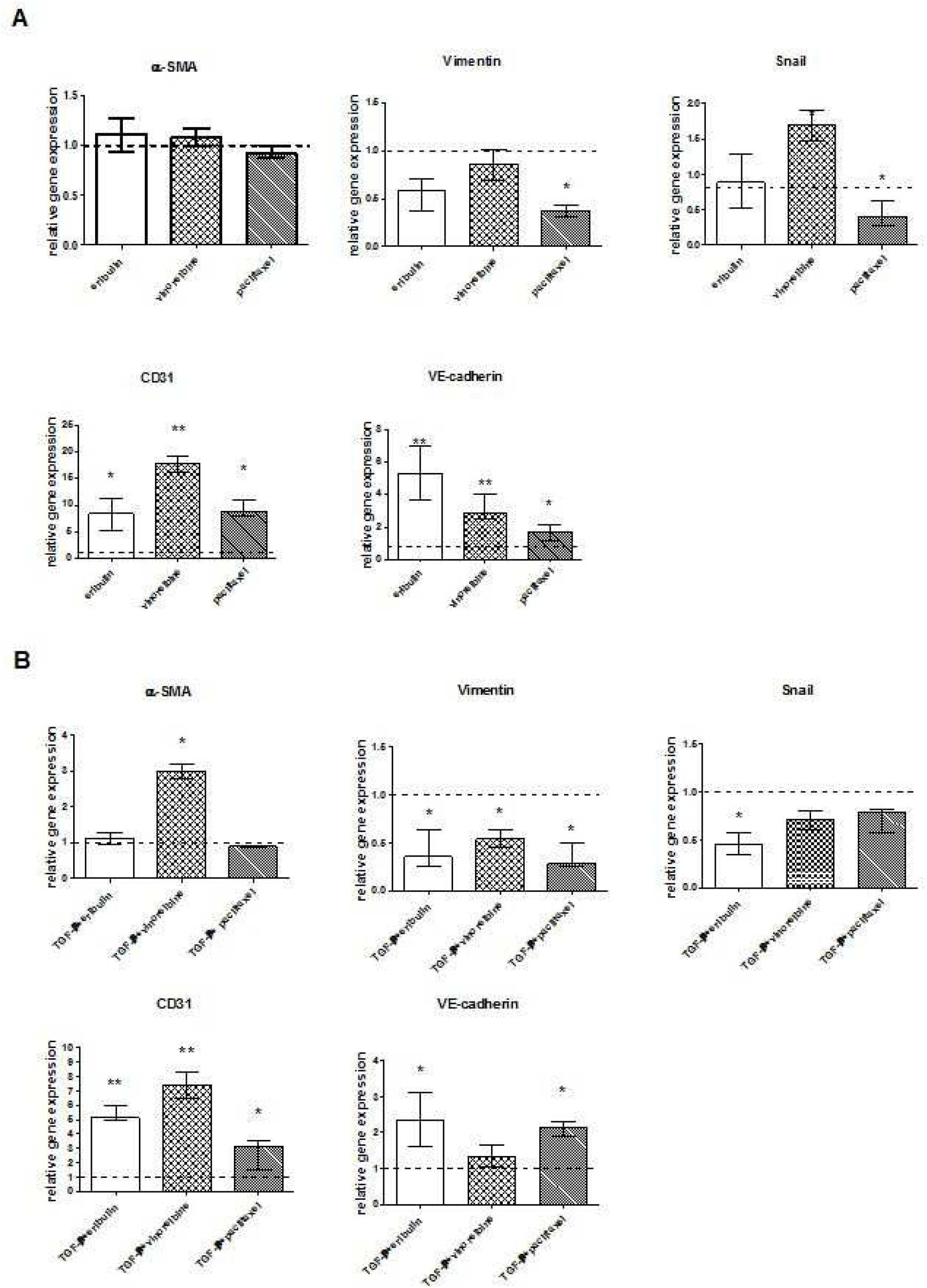
Analysis of the transcription of mesenchymal and epithelial molecular markers in HUVEC treated with drugs targeting the microtubule system. (A) Transcripts were measured by qPCR analysis in HUVEC after 4 hrs of treatment with either 1.0 nM eribulin, 0.8 nM paclitaxel, 10 nM vinorelbine or medium (CTR). (B) qPCR analysis was done after 4 hrs of cell pre-incubation with 10 ng/mL TGF-β and 4 more hours of drug treatment. Bars represent data obtained from the mean of four independent experiments (± 1 SD); samples were normalized to CTR represented as the dashed line at 1. (** p<0.01, *p<0.05)

TGF-β increases the transcription of the three mesenchymal markers and decreases the transcription of the two epithelial markers (15).

We assessed how the three drugs affected TGF-β modulation of the EndMT markers by checking the effect on HUVEC pre-incubated with TGF-β.

All treatments showed a significant down-regulation of Vimentin mRNA compared to CTR.

The transcription of αSMA was significantly increased by vinorelbine, decreased by paclitaxel and unaffected by eribulin. The expression of Snail was significantly down-regulated by eribulin (Fig. 2B).

CD31 mRNA was significantly enhanced by all drugs, in particular by eribulin and vinorelbine, and to a lesser extent by paclitaxel. Eribulin and paclitaxel significantly increased the expression of VE-cadherin (Fig. 2B).

To evaluate the overall effect of the drugs on both epithelial and mesenchymal markers, we calculated a *ratio* between the expression of epithelial and mesenchymal markers (E/M). Compared to CTR, a higher E/M *ratio* was found for each of the three drugs (Fig 3A). However, compared to paclitaxel and vinorelbine, eribulin better counteracted TGF-β-induced EndMT, since the E/M *ratio* on TGF-β pre-treated HUVEC was significantly higher compared to the *ratio* of the other two drugs (Fig. 3B).

**Figure 3.**
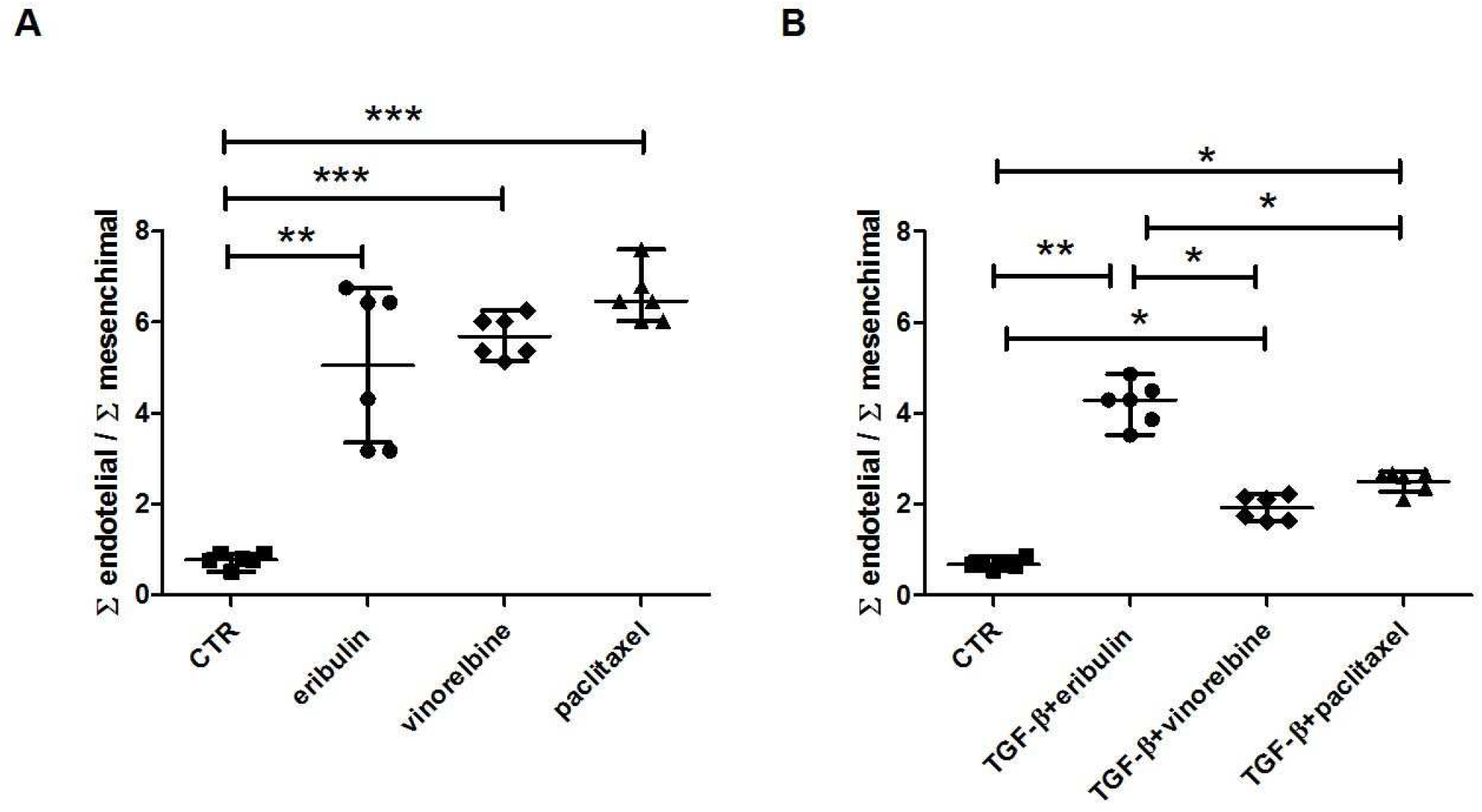
Analysis of the *ratio* between epithelial and mesenchymal markers. The *ratio* was calculated between the sum of the *ratio* of the epithelial markers and the sum of the *ratio* of the mesenchymal markers (E/M). (A) Expression analysis after 4 hr of drug treatment. (B) Expression analysis after 4 hr of pre-treatment with TGF-β and 4 more hours of drug treatment. The *ratio* is expressed on the ordinates as a value derived from the exponential in base two of the ΔΔCT (2^-ΔΔCt^). (*** p<0.001, ** p<0.01, * p<0.05)

### 3.3 Eribulin increases ICAM-1 expression on HUVEC

Finally, we evaluated the effect of eribulin on the expression of ICAM-1 and VCAM-1 adhesion molecules, involved in the process of leukocyte adhesion, rolling, and extravasation (16).

In CTR, VCAM-1 was undetectable while ICAM-1 was expressed at low levels (Fig. 4).

**Figure 4.**
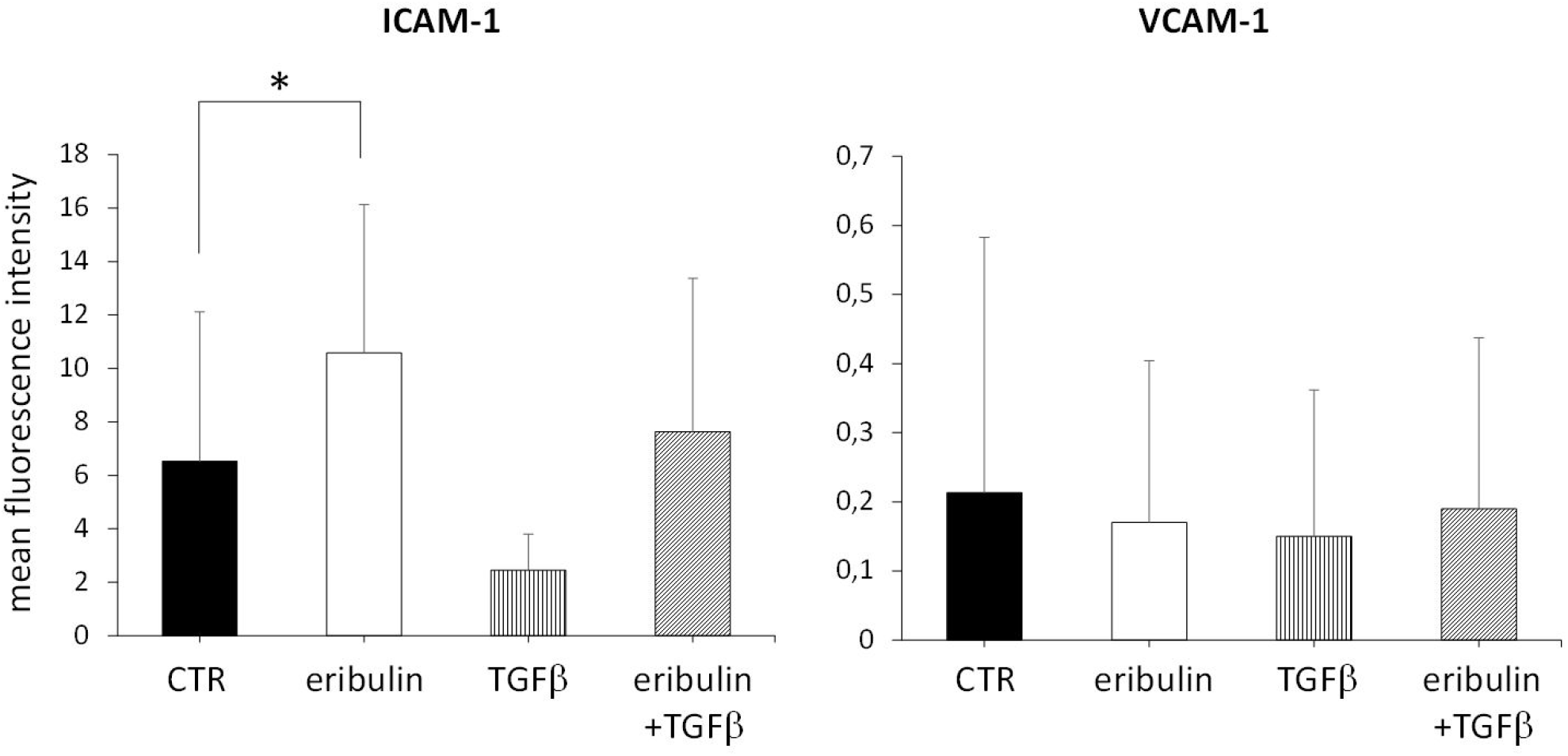
Flow cytometry analysis of ICAM-1 and VCAM-1 expression in HUVEC treated with eribulin and/or TGF-β. Cells were incubated for 24 hrs with medium (CTR), 1 (VCAM-1) or 10 (ICAM-1) ng/mL TGF-β and/or 5 nM eribulin. Mean + 1 SD of three independent experiments. * P<0.05.

The inducibility of these markers was preliminarily checked by incubating the HUVEC with TNF-α which greatly increased the expression of both ICAM-1 and VCAM-1 (Supplementary Fig.1). Adding eribulin to CTR significantly increased the expression of ICAM-1 without any effect on VCAM-1.

TGF-β is known to downregulate ICAM-1 and VCAM-1 (17)(18).

We checked the effect of eribulin on ICAM-1 and VCAM-1 expression modulated by TGF-β.

Eribulin was able to counteract the downregulation of ICAM-1 induced by TGF-β (Fig. 4).

## 4 Discussion

In our study eribulin exerted an antiangiogenic activity on human endothelial cells. Similar effect was previously observed by Agoulnik et al (7). In particular, we found that eribulin displays a dual antiangiogenic effect: i) it prevents the organization of the vascular reticulum during the first step of morphogenesis, and ii) it is also able to counteract the EndMT effect mediated by TGF-β.

Eribulin, exerts its therapeutic effect primarily by targeting the microtubule cytoskeleton, however off-target effects have been reported and could have important implications in its therapeutic activity.

We herein demonstrated that eribulin can affect the endothelial cell organization by interfering with the morphogenesis process. At the molecular level the drug did not impact the transcription of any of the mesenchymal markers. However, it increases the transcription of the epithelial markers CD31 and VE-cadherin in both untreated and TGF-β-treated cells. Both CD31 and VE-cadherin are instrumental in endothelial cell organization and function by reshaping and strengthening the endothelial lining and by controlling vascular permeability (19). Our results provide the molecular basis to defining that eribulin treatment induces vascular remodeling associated with improved perfusion in breast cancer xenograft models (2).

CD31 and VE-cadherin are also instrumental in controlling leukocyte extravasation (19, 20). In particular CD31 plays a fundamental role in the extravasation not only myeloid and natural killer (NK) cells but of T-cells as well (21). Eribulin treatment also significantly increased ICAM-1 membrane expression in CTR and shows a clear trend toward the downregulation of the molecule induced by TGF-β. ICAM-1 is downregulated in tumor-derived endothelial cells (22) and, since ICAM-1 is involved in leukocyte adhesion and endothelial cells extravasation (23), this phenomenon may also contribute to dampen the recruitment of T and NK cells into the tumors. The findings mentioned above supported the hypothesis that eribulin may in part exerts its pharmacological effect by promoting tumor infiltration by immune effector cells through the induction of the overexpression in endothelia of molecules involved in leukocyte extravasation. Indeed, in a xenograft tumor model, eribulin treatment was found to increase the number of tumorinfiltrating immune cells, and NK cells depletion reduced the antitumor effects of eribulin (24).

These effects of eribulin on immune cells recruitment at the tumor site is important, since many of the human solid tumors, including breast cancer, show an excluded immunophenotype (4, 25).

We gathered evidence that eribulin may trigger another therapeutic mechanism. The induction of EMT by TGF-β is well known in literature, and several studies indicated increased TGF-β signaling as a key effector of EMT in cancer progression and metastasis (26, 27).

However, TGF-β has been found to play a central role also in inducing EndMT (28, 29), not surprisingly since EMT and EndMT are supported by almost identical mechanism. TGF-β induces EndMT via Smad signaling, which results, among other effects, in increased transcription of Snail, αSMA and Vimentin, and downregulation of VE-cadherin and CD31 (30). Eribulin significantly inhibited the TGF-β mediated effect on the transcription of Snail, Vimentin, VE-cadherin and CD31, suggesting the part of the pharmacological effect of the drug is the result of its ability to inhibit the induction of EndMT by TGF-β. Interestingly, we had previously observed that response to eribulin in breast cancer patients correlated to a reduction of circulating TGF-β (3).

On the basis of our results, we suggest that eribulin exerts its pharmacological effect also via three off target mechanisms: i) increases of transcription of CD31 and VE-cadherin, which results in vascular remodeling, ii) promotes tumor infiltration by immune cells by increasing transcription of CD31 and VE-cadherin, and expression of ICAM-1, and iii) counteracts the EndMT mediated by TGF-β. These conclusions might indicate new therapeutic approaches. For instance, we could speculate that eribulin might achieve the best therapeutic effect in tumors displaying an immune excluded phenotype.

Vinorelbine and paclitaxel are known to exert antiangiogenic activity on HUVEC (13), however we found that the transcription patterns are different between them and compared to eribulin. Vinorelbine upregulated the transcription of Snail, CD31 and VE-cadherin in CTR, and of αSMA, Vimentin and CD31 in HUVEC pre-treated with TGF-β.

Paclitaxel reduced the transcription of Vimentin and Snail in CTR and of Vimentin and αSMA in TGF-β-treated cells, but increased the transcription of the epithelial markers in both treated and untreated HUVEC. These findings suggest that each of the three drugs might affect endothelial cell function through distinctive mechanisms. When the individual gene expressions were added together in macro-groups, generating a *ratio* between endothelial and mesenchymal characters, eribulin showed the same *ratio* of the other drugs on CTR, but a higher *ratio* on TGF-β-treated cells. This finding strengthens our hypothesis on the role of eribulin in EndMT, confirming that its activity takes place also by counteracting the TGF-β-dependent polarization of the endothelial cells towards the mesenchymal phenotype.

TGF-β is highly expressed in many solid tumors and in particular in excluded tumors which are characterized by a TGF-β signature (5).

Many human breast cancers show an excluded immunophenotype (6) and interestingly, we have observed that response to eribulin in breast cancer correlates to a reduction of circulating TGF-β (3).

Therefore, considering the off-target effects of eribulin, this drug might favor the immune checkpoint inhibitors (ICIs) activity in excluded breast cancer, and possibly in other excluded solid tumors, by removing many mechanisms of T-cell exclusion.

## Author Contributions

A.A., A.F., M.P., S.A., and S.M. planned and carried out the experiments. M.P. performed also data analysis. N.D and F.R. contributed to the drafting of manuscript. OB., M.C.M and O.G. were involved in planning, supervised and wrote the work. A.A. and A.F. contributed equally to the work and share first authorship. M.C.M and O.G. contributed equally to the work and share last authorship.

## References

1. Garrone, O., G., Miraglio, E., Vandone A.M., Vanella, P., Lingua, D., Merlano, M.C., (2017). Future oncology. 13(30):2759–2769. doi: 10.2217/fon-2017-0283.

2. Funahashi, Y., Okamoto, K., Adachi, Y., Semba, T., Uesugi, M., Ozawa, Y. et al. (2014). Eribulin mesylate reduces tumor microenvironment abnormality by vascular remodeling in preclinical human breast cancer models. Cancer Sci. 105(10):1334–42.

3. Garrone, O., Michelotti, A., Paccagnella, M., Montemurro, F., Vandone, A.M., Abbona, A. et al. (2020). Exploratory analysis of circulating cytokines in patients with metastatic breast cancer treated with eribulin: the TRANSERI-GONO (Gruppo Oncologico del Nord-Ovest) study. ESMO Open. 5:e000876. doi:10.1136/esmoopen-2020-000876

4. Ueda, S., Saeki, T., Takeuchi, H. Shigekawa, T., Yamane, T., Kuji, I., et al. (2016). In vivo imaging of eribulin-induced reoxygenation in advanced breast cancer patients: a comparison to bevacizumab. Br. J. Cancer. 114:1212–8. doi: 10.1038/bjc.2016.122

5. Hedge, P.S. and Chen, D.S. (2020). Top 10 challenges in cancer immunotherapy. Immunity. 52:17–35. doi: 10.1016/j.immuni.2019.12.011.

6. Bray, F., Ferlay, J., Soerjomatam, I., Siegel, R.L., Torre, L.A., Jernal, A. (2018). Global cancer statistics 2018: GLOBOCAN estimates of incidence and mortality worldwide for 36 cancers in 185 conuntries. CA Cancer J. Clin. 68, 394–424. doi: 10.3322/caac.21492.

7. Agoulnik, S.I., Kawano, S., Taylor, N., Oestreicher, J., Matsui, J., Chow, J., et al. (2014). Eribulin mesylate exerts specific gene expression changes in pericytes and shortens pericyte-driven capillary network in vitro. Vasc Cell 6, 3. doi: 10.1186/2045-824X-6-3

8. Fontani, F., Domazetovic, V., Marcucci, T., Vincenzini M.T., Iantomasi, T. (2016). Tumor Necrosis Factor-Alpha Up-Regulates ICAM-1 Expression and Release in Intestinal Myofibroblasts by Redox-Dependent and -Independent Mechanisms. J. Cell Biochem. 117:370–381. doi: 10.1002/jcb.25279

9. Livak, K. J., and Schmittgen T. D. (2001). Analysis of relative gene expression data using realtime quantitative PCR and the 2^-△△CT^ method. Methods. 25, 402–408. doi: 10.1006/meth.2001.1262.

10. Piera-Velazquez, S., Li, Z., Jimenez, S.A. (2011). Role of endothelial-mesenchymal transition (EndoMT) in the pathogenesis of fibrotic disorders. Am. J. Pathol.;179:1074–1080. doi: 10.1016/j.ajpath.2011.06.001.

11. Crabtree, B., and Subramanian, V. (2007). Behavior of endothelial cells on Matrigel and development of a method for a rapid and reproducible in vitro angiogenesis assay. In Vitro Cell Dev Biol Anim. 43(2):87–94. doi: 10.1007/s11626-007-9012-x.

12. Piera-Velazquez, S., and Jimenez, S.A. (2019). Endothelial to Mesenchymal Transition: Role in Physiology and in the Pathogenesis of Human Diseases. Physiol Rev. 99(2):1281–1324. doi: 10.1152/physrev.00021.2018.

13. Mavroeidis, L., Sheldon, H., Briasoulis, E., Marselos, M., Pappas, P., Harris, A.L. (2015). Metronomic vinorelbine: Anti-angiogenic activity in vitro in normoxic and severe hypoxic conditions, and severe hypoxia-induced resistance to its anti proliferative effect with reversal by Akt inhibition. Int J Oncol. 47(2): 455–464. doi: 10.3892/ijo.2015.3059.

14. Bocci, G., Di Paolo, A., Danesi, R. (2013). The pharmacological bases of the antiangiogenic activity of paclitaxel. Angiogenesis. 16(3):481–92. doi: 10.1007/s10456-013-9334-0.

15. Kalluri, R., and Weinberg, R.A. (2009). The basics of epithelial-mesenchymal transition. J Clin Invest. 119(6):1420–8. doi: 10.1172/JCI39104.

16. Vestweber, D. (2015). How leukocytes cross the vascular endothelium. Nat Rev Immunol. 15(11):692–704. doi: 10.1038/nri3908.

17. Kiyohara, H., Ishizaki, Y., Suzuki, Y., Katoh, H., Hamada, N., Ohno, T., et al (2011). Radiation-induced ICAM-1 expression via TGF-β1 pathway on human umbilical vein endothelial cells; comparison between X-ray and carbon-ion beam irradiation. J Radiat Res. 52(3):287–92. doi: 10.1269/jrr.10061.

18. Gamble, J.R., Bradley, S., Noack, L., Vadas, M.A. (1995). TGF-beta and endothelial cells inhibit VCAM-1 expression on human vascular smooth muscle cells. Arterioscler Thromb Vasc Biol. 15(7):949–55. doi: 10.1161/01.atv.15.7.949.

19. Bravi, L., Dejana, E., Lampugnani, M.G. (2014). VE-cadherin at a glance. Cell Tissue Res. 355, 515–522. doi:10.1007/s00441-014-1843-7.

20. Schimmenti, L.A., Yan, H.C., Madri, J.A., Albelda, S.M. (1992). Platelet endothelial cell adhesion molecule, PECAM-1, modulates cell migration. J Cell Physiol. 153(2):417–28. doi: 10.1002/jcp.1041530222.

21. Marelli-Berg, F.M., Clement, M., Mauro, C., Caligiuri, G. (2013). An immunologist’s guide to CD31 function in T-cells. J Cell Sci. 126(Pt 11):2343–52. doi: 10.1242/jcs.124099.

22. Griffioen, A.W., Damen, C.A., Blijham, G.H., Groenewegen, G. (1996). Tumor angiogenesis is accompanied by a decreased inflammatory response of tumor-associated endothelium. Blood. 88(2):667–73. PMID: 8695814.

23. Schnoor, M., Alcaide, P., Voisin, M.B., van Buul, J.D. (2015). Crossing the Vascular Wall: Common and Unique Mechanisms Exploited by Different Leukocyte Subsets during Extravasation. Mediators Inflamm.;2015:946509. doi: 10.1155/2015/946509.

24. Ito, K., Hamamichi, S., Abe, T., Akagi, T., Shirota, H., Kawano, S., et al. (2017). Antitumor effects of eribulin depend on modulation of the tumor microenvironment by vascular remodeling in mouse models. Cancer Sci. 108(11):2273–2280. doi: 10.1111/cas.13392.

25. Goldberg, J., Pastorello, R.G., Vallius, T., Davis, J., Cui, Y.X., Agudo, J., et al. (2021). The Immunology of Hormone Receptor Positive Breast Cancer. Front Immunol. 12:674192. doi: 10.3389/fimmu.2021.674192.

26. Wendt, M.K., Allington, T.M., Schiemann, W.P. (2009). Mechanisms of the epithelial-mesenchymal transition by TGF-beta. Future Oncol. 5(8):1145–68. doi: 10.2217/fon.09.90.

27. Dudas, J., Ladanyi, A., Ingruber, J., Steinbichler, T.B., Riechelmann, H. (2020). Epithelial to Mesenchymal Transition: A Mechanism that Fuels Cancer Radio/Chemoresistance. Cells. 9(2):428. doi: 10.3390/cells9020428.

28. Medici, D., Shore, E.M., Lounev, V.Y., Kaplan, F.S., Kalluri, R., Olsen, B.R (2010). Conversion of vascular endothelial cells into multipotent stem-like cells. Nat Med. 16(12):1400–6. doi: 10.1038/nm.2252.

29. Yoshimatsu, Y., Watabe, T. (2011). Roles of TGF-β signals in endothelial-mesenchymal transition during cardiac fibrosis. Int J Inflam 2011:724080. doi: 10.4061/2011/724080.

30. Ma J, Sanchez-Duffhues G, Goumans MJ, Ten Dijke P. TGF-β-Induced Endothelial to Mesenchymal Transition in Disease and Tissue Engineering. Front Cell Dev Biol. 2020 Apr 21;8:260. doi: 10.3389/fcell.2020.00260.

