## supplementary Table 1 for "Effect of eribulin on angiogenesis and endothelial adhesion molecules"

### *Supplementary Material*

**Supplementary Table 1 - Gene analyzed and primers used.** Primer sequences and melting temperature ($T_M$) of each primer used in the present study are listed.

| Gene name | Primer sequences | Melting temperature $T_M$ (°C) |
|---|---|---|
| CD31 | LEFT PRIMER: GTCAGCAGCATCGTGGTCAAC | 64.5 |
| | RIGHT PRIMER: CCACTGTCCGACTTTGAGGC | 64.5 |
| Vimentin | LEFT PRIMER: CACCTGTGAAGTGGATGCC | 62.3 |
| | RIGHT PRIMER: AATCCTGCTCTCCTCGCCTT | 62.45 |
| α-SMA | LEFT PRIMER: ACCCTGTTCCAGCCATCCTT | 62.45 |
| | RIGHT PRIMER: TTGCGGTGGACAATGGAAGG | 62.45 |
| VE-Cadherin | LEFT PRIMER: CAAGCCCTACCAGCCCAAAG | 64.5 |
| | RIGHT PRIMER: CCCGGTCAAACTGCCCATAC | 64.5 |
| Snail1 | LEFT PRIMER: CTATGCCGCGCTCTTTCCTC | 60.6 |
| | RIGHT PRIMER: AGGAGAAGGACGAAGGAGCC | 58.8 |
| G6PDH | LEFT PRIMER: ACAACGATACCAGGGTGTTACC | 62.35 |
| | RIGHT PRIMER: TCTCCCATGATGCCTTTAACTC | 62.3 |

**Supplementary Figure 1 – Flow cytometry evaluation of the inducibiliy of ICAM-1 and VCAM-1 expression in HUVEC.** Cells were incubated for 24 hrs with medium (CTR) or 10 ng/mL TNF-α. Mean +1 SD of three independent experiments.

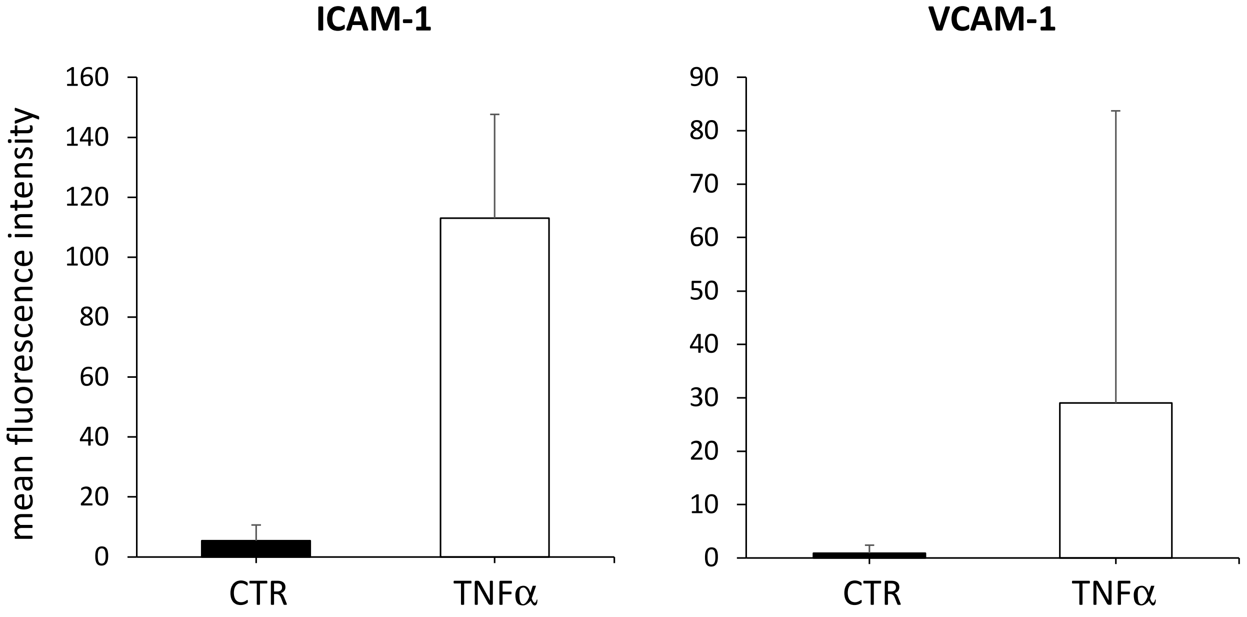
